# Fast Vesicle Fusion Mediated by Hydrophobic Dipeptides

**DOI:** 10.1101/2020.11.19.390179

**Authors:** C. Wei, A. Pohorille

## Abstract

To understand the transition from inanimate matter to life, we studied a process that directly couples simple metabolism to evolution via natural selection, demonstrated experimentally by Adamala and Szostak (Nat. Chem. 2013, 5, 495–501). In this process, dipeptides synthesized inside precursors of cells promote absorption of fatty acid micelles to vesicles inducing their preferential growth and division at the expense of other vesicles. The process is explained on the basis of coarse-grained molecular dynamics simulations, each extending for tens of microseconds, carried out to model fusion between a micelle and a membrane, both made of fatty acids in the absence and presence of hydrophobic dipeptides. In all systems with dipeptides, but not in their absence, fusion events were observed. They involve the formation of a stalk made by hydrophobic chains from the micelle and the membrane, similar to that postulated for vesicle-vesicle fusion. The emergence of a stalk is facilitated by transient clusters of dipeptides, side chains of which formed hydrophobic patches at the membrane surface. Committor probability calculations indicate that the size of a patch is a suitable reaction coordinate and allow for identifying the transition state for fusion. Free energy barrier to fusion is greatly reduced in the presence of dipeptides to only 4-5 kcal/mol, depending of the hydrophobicity of side chains. The mechanism of mediated fusion, which is expected to apply to other small peptides and hydrophobic molecules, provides a robust means by which a nascent metabolism can confer evolutionary advantage to precursors of cells.

**Significance:** We study fusion of micelles and vesicles in the presence and absence of hydrophobic dipeptides by way of molecular dynamics simulations and demonstrate the spontaneous formation of a hydrophobic stalk that has been also long postulated as the key process in vesicle-vesicle fusion but shown only in forced simulations. We show that fusion is facilitated by clusters of dipeptides that form hydrophobic patches at the membrane surface. In order to understand the transition from inanimate matter to life, we explain and generalize experimental findings that hydrophobic dipeptides synthesized inside precursors of cells promote absorption of fatty acid micelles to vesicles inducing their preferential growth and division, thus providing cells endowed with such metabolism with evolutionary advantage.

## Introduction

How inanimate matter transitioned to animate structures remains a major, unanswered question at the intersection of chemistry and biology. Identifying processes that might have contributed to this transition would not only advance understanding of the origins of life (1–4), but also drive evolutionary approaches to synthetic biology and, in particular, to building artificial cells (2, 5–8). In this paper, we apply molecular dynamics computer simulations to elucidate the general mechanism of a process that directly links specific chemical reactions in a nascent metabolism to the growth and division of ancestors of cells, thus providing them with advantage in evolution through natural selection.

It is broadly accepted that modern cells were preceded by protocells (2, 9–12) – self-assembled, membrane bound structures that encapsulated the emerging metabolism and information carriers. Considering that phospholipids were unlikely to be present in the prebiotic milieu (11, 13, 14), it has been postulated that ancestral membranes were formed by heterogeneous amphiphilic material containing single hydrocarbon chain species, presence of which was plausible on the early Earth (12, 14–17). Commonly used laboratory models for envelopes of the earliest protocells are vesicles formed by fatty acids (18–26).

The content of the earliest protocells was most likely highly heterogeneous. This set the stage for evolution by natural selection (1). Competing for limited resources, populations of protocells capable of producing more environmentally fit progeny would increase in time at the expense of other protocells. A simple mechanism to achieve such competitive advantage would be to encapsulate metabolism that promotes growth and division of vesicles. Processes that could induce competitive growth and subsequent division include synthesis of membrane forming material from its precursors (23), stressing vesicles osmotically with encapsulated replicator of nucleic acids (27), photochemically driven redox chemistry (28), regulation of catalytic activity (29), stabilization of the protocellular compartment (26) or the incorporation of small amounts of phospholipids into vesicle membrane (30).

A potentially highly significant mechanism that directly links simple metabolism to vesicle growth and division has been identified by Adamala and Szostak (24). Seryl-histidine dipeptides and two terminally blocked amino acids, N-acetyl-L-phenylalanine ethyl ester and leucinamide were encapsulated in oleic acid vesicles. The dipeptide, which was shown to act as a weak catalyst of peptide bond formation (31–34), mediated the synthesis of a dipeptide, N-acetyl-L-phenylalanine leucinamide (Phe-Leu), from the amino acid substrates. The presence of this newly synthesized, highly hydrophobic dipeptide drove vesicle growth when excess fatty acids in the form of individual molecules or micelles were present in the medium. Vesicles containing Phe-Leu grew at the expense of those lacking the dipeptide if both were incubated together. As they became larger, they adapted fragile, filamentous structures that could easily fragment to produce offspring vesicles (28, 35).

Although it was established that peptide hydrophobicity was important for vesicle growth, neither the underlying mechanism nor the extent to which this mechanism applied to other hydrophobic peptides and metabolites were determined. Identifying this mechanism, as it emerges from molecular dynamics computer simulations, is the central theme of this paper. It builds on our previous work in which we have shown that hydrophobic, but not hydrophilic, dipeptides tend to accumulate at the water-vesicle interface and that this tendency is correlated with the degree of hydrophobicity (36).

In this paper, we concentrate on the key step in the process observed by Adamala and Szostak (24) - fusion of vesicles with micelles in the absence and presence of different dipeptides at the surface of the membrane. This type of fusion is poorly understood. A computational study on antimicrobial lipopeptides in which micelles formed by a cationic tetrapeptide attached to palmitoyl chain were inserted into phospholipid membranes revealed significant barriers to this process that depended on the composition of the membrane (37). In this study, we demonstrate a different underlying mechanism that is remarkably similar to the first, key step in vesicle-vesicle fusion (38–40). In both types of fusion, this step is the formation of a stalk joining the two structures. We further show that the presence of nonpolar dipeptides facilitates this process through the formation of hydrophobic patches due to the clustering of the hydrophobic amino-acid side chains at the surface of the membrane. This, in turn, markedly decreases the free energy barrier to fusion. The ability to form hydrophobic patches does not appear to be limited to dipeptides, but most likely applies to a variety of molecules that contain hydrophobic groups. This points to a general mechanism in which synthesis of such molecules conferred evolutionary advantage in the earliest stages of protocellular life.

## Methods

Fusion processes involving vesicles occur at sub-millisecond to second time scales (41, 42). Due to computational costs, this is an obstacle for all atom molecular dynamics (AAMD) simulations. For this reason, we used instead coarse-grained molecular dynamics (CGMD), as it can markedly accelerate simulations. In CGMD, several atoms are grouped into a single bead, which not only significantly reduces the number of particles in the system, but also has a smoothing effect on rugged free energy landscapes. In this study, we used the MARTINI force field (43, 44), as implemented in the NAMD package (45). This force field has been already widely applied to understand cell membranes (46) vesicle formation (47) and membrane or vesicle fusion (40, 48, 49).

CGMD simulations were carried out in a cell, the x, y, z dimensions of which were approximately 95×95×190 Å. The z-dimension of the cell was perpendicular to the membrane surface. Periodic boundary conditions were applied in all three directions. The system contained a planar membrane with embedded blocked dipeptides and a micelle, all dissolved in water. Both the membrane bilayer and the micelle consisted of oleic acid (OA), as in the experimental study of Adamala and Szostak (24). There were 19924 water molecules and 612 OA molecules in the system, of which 100 OA molecules formed the micelle. This is the experimentally determined aggregation number for OA micelles (50). 318 Na^+^ and 62 Cl^−^ were also included in the unit cell, which corresponds to an ionic concentration of 0.2 M (excluding cations required to neutralize the system).

pK_a_ of OA in aqueous solution at 25° C is 5.0 (51). In the membrane, its value increases markedly to approximately 8.0 – 8.5 (52–54). An increase to 9.85 was also observed in OA monolayers spread on water (55). This is due to cooperative effects between two adjacent carboxyl groups, which can form strong hydrogen bonds if one is in a protonated form and the other is deprotonated (16, 56). In micelles, however, spacing between head groups of neighboring OA molecules is markedly larger and, therefore, the formation of such hydrogen bonds is much less likely. As a consequence, pK_a_ of OA in micelles is close to that in aqueous solution (57). To reproduce experimental conditions of nearly neutral pH (24) in the simulations, the ratio of protonated and deprotonated OA molecules in the membrane was set to 1:1, whereas all OA molecules were deprotonated in the micelle. The surface area per OA head group in the membrane was 35 Å^2^, which is close to the surface area of 33 Å^2^ obtained from NMR data at a temperature of 303 K (52), which is slightly lower than 310 K applied in our simulations.

The Phe-Leu dipeptide and other three dipeptides studied here, composed of two phenylalanines (Phe-Phe), valines (Val-Val) and leucines (Leu-Leu), were blocked with the acetyl group at the N-terminal and N-methylamide at the C-terminal. The latter differs from the amide blocking group used in the experimental work (24) because the MARTINI force field lacks parametrization for amide at the C-terminus of a peptides. The impact of this difference on fusion mechanism is expected to be negligible. For each simulation, 50 dipeptides were initially embedded evenly in the two leaflets of the membrane, yielding a molar ratio of 10% with respect to OA molecules.

The simulations were carried out in NPT ensemble with pressure of 1 bar controlled by way of Nosé-Hoover Langevin pressure piston. Temperature of 310 K was kept constant by way of Langevin dynamics with a friction damping coefficient of 5 ps^−1^. Time step of 20 fs suggested for CGMD-MARTINI simulations with proteins (58) was used to integrate Newton’s equations of motion. Long-ranged electrostatic interactions were handled by way of particle mesh Ewald (PME) method with a grid of 100×100×200, together with a dielectric constant of 15 (58–60). The length of the CGMD trajectories in the vesicle/micelle/peptide systems varied from 1 to 20 *μs*. Each trajectory was terminated once an irreversible step in fusion, as described further in the text, had occurred.

To understand adsorption of micelle onto the fatty acid membrane, free energy profile, *A(z)*, was calculated with the aid of the Adaptive Biasing Force (ABF) method (61, 62) in which the reaction coordinate, *z*, was chosen as the distance between the z-components of the center mass of the micelle and the center of mass of the membrane. The range of 46 Å < *z* < 64 Å, which corresponds to the transfer of the micelle from the membrane surface to the water phase, was divided into three windows (46 Å < *z* < 52 Å, 52 Å < *z* < 58 Å, and 58 Å < *z* < 64 Å). The micelle was restrained to remain within each window with a harmonic potential acting from the edges of the window outwards. For each window, an ABF simulation 800 ns in length was carried out.

To validate CGMD with the MARTINI force field, results of coarse-grained and AAMD simulations for the OA membrane with and without dipeptides were compared. In AAMD, the updated version of CHARMM potentials for phospholipids (63, 64) and the recently optimized CHARMM force field for proteins (65) were used. Water was represented by way of TIP3P model (66). In Fig. S1, density profiles of different components in the pure OA membrane system are shown. The distributions of OA (both tails and head group), water, and ions obtained from all-atom and coarse-grained approaches were found to be in good agreement, although several small differences, such as a slightly deeper penetration of water into the membrane in AAMD, were observed. The peak in the density of Na^+^ at the membrane surface, which is due to electrostatic attraction between sodium cations and negatively charged, deprotonated OA molecules in the membrane, was found in both simulations, although the peak was sharper in the CGMD simulation. In this simulation, but not in AAMD, there was also a small interfacial peak in the density of Cl^−^, but its effect is expected to be negligible, compared to interactions between negatively charged membrane and cations. Further, CGMD and AAMD simulations of 3 *μs* and 300 *ns* in length, respectively, were carried out for the OA membrane with 50 molecules of the Phe-Leu dipeptide. The free energy profiles, *A(z)*, calculated from the density distribution of the dipeptide along the membrane normal, *p(z)*, according to the Boltzmann distribution, 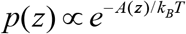, is shown in Fig. S2. The agreement between *A(z)* obtained from CGMD and AAMD is good. In both cases, favorable adsorption of the dipeptide at the water-membrane interface and a barrier of 3 - 4 kcal/mol at the center of the membrane were found. Although both side chains of the dipeptide are hydrophobic, the polar backbone makes the molecule amphiphilic, which accounts for its tendency to accumulate at the interface. In general, a consistent agreement between CGMD-MARTINI and AAMD-CHARMM simulations of the OA membrane with and without a dipeptide suggests that CGMD is a suitable simulation tool for these and related systems.

## Results and Discussion

The mechanism of fusion between a micelle and an OA membrane in the absence and presence of dipeptides was studied by way of CGMD simulations at time scales extending for tens of microsecond. Besides the Phe-Leu dipeptide that was investigated experimentally by Adamala and Szostak (24), three other hydrophobic dipeptides, Phe-Phe, Val-Val, and Leu-Leu, were also embedded in the membrane and studied for their effects on fusion. All dipeptides were evenly distributed between the two leaflets of the membrane. Even though the hydrophobic peptides are generated experimentally inside a vesicle (24), using the even distribution was justified because the free barrier to permeation of hydrophobic dipeptides across the membrane, approximately equal to 3 - 4 kcal/mol (36), is sufficiently low that they equilibrate on a time scale faster than the time scale of fusion. In all simulations, the micelle was found to start fusing with the membrane in the presence of dipeptides in several to tens of μs following a general mechanism, which always involved the formation of a stalk structure. We first present results for simulations without dipeptides, next we focus on the experimentally investigated Phe-Leu system, and subsequently discuss the other three dipeptide systems.

### Micelle-membrane interactions in the absence of peptides

Direct interactions between the OA bilayer and an approaching micelle are repulsive, as both are negatively charged. These interactions, however, are significantly modulated by positively charged sodium ions near the surfaces of both the bilayer and the micelle. As a result, the association is energetically favored. This can be seen in Fig. S3, which depicts the free energy profile, *A(z)*, characterizing bilayer-micelle interactions. The free energy decreases monotonically by 4.8 kcal/mol over a range of approximately 14 Å and reaches the minimum at contact. This means that association of a micelle and a vesicle is spontaneous and barrier less. In subsequent simulations of the micelle initially set to be in contact with the bilayer, which extended for 90 *μs*, the micelle remained at the surface of the membrane. No signs of fusion or micelle dissociation from the surface of the vesicle were observed. The former outcome is consistent with the slow timescales of vesicle fusion (41, 42). The latter result is expected considering that the calculated diffusion coefficient of micelles in water is quite small, of the order of 10^−8^ cm^2^/s. This means that for the calculated interfacial minimum the dissociation time should be on the order of milliseconds.

### Fusion of micelle with OA membrane: stalk formation mediated by dipeptides

The behavior of the membrane-micelle complex in the presence and absence of dipeptides differed considerably in the CGMD simulations. In contrast to the pure complex, which did not exhibit signs of fusion in a 90 *μs* simulation, the micelle was found to fuse with the membrane containing Phe-Leu through an hourglass shaped stalk structure that formed within 1.2 to 12.2 *μs* in eight independent simulations.

Since fusion can be regarded as mixing of OA molecules from two initially separate reservoirs, it can be conveniently tracked by way of a parameter, *N*_*t*_, defined as the total number of tail beads in OA molecules forming the membrane that are within the van der Waals distance of 5.27 Å from any tail bead of OA molecules in the micelle. This distance corresponds to the minimum in the Lennard-Jones potential between two beads in the MARTINI force field. As shown in Fig. 1a for Phe-Leu, *N*_*t*_ was zero for most of the first 6.7 *μs* of the trajectory. This indicated that there was no contact between the hydrophobic tails of OA molecules from the micelle and the membrane. A representative snapshot of such structure of the complex is shown in Fig. 1b. Only occasionally, a few beads from the micelle and the membrane intermittently came in contact. This happened at times, *t*, equal to 1.2, 2, and 2.2 *μs*, as marked in Fig. 1a. These events can be interpreted as early, unsuccessful fusion attempts. After 6.7 *μs*, an abrupt jump in *N*_*t*_ was observed, whereby hydrophobic tails of OA from the membrane and the micelle began to mix in the contact region. This signaled the beginning of fusion. A cluster of Phe-Leu dipeptides was also present in the contact area, as shown in the snapshot from the simulation, shown in Fig. 1c. Within tens of nanoseconds, a stalk joining the micelle and the membrane formed in which the hydrophobic tails of fatty acids occupied the contact area between the membrane and the micelle, whereas ions and polar groups are displaced to the boundary of the contact zone. This is shown in Fig. 1d.

**Figure 1.**
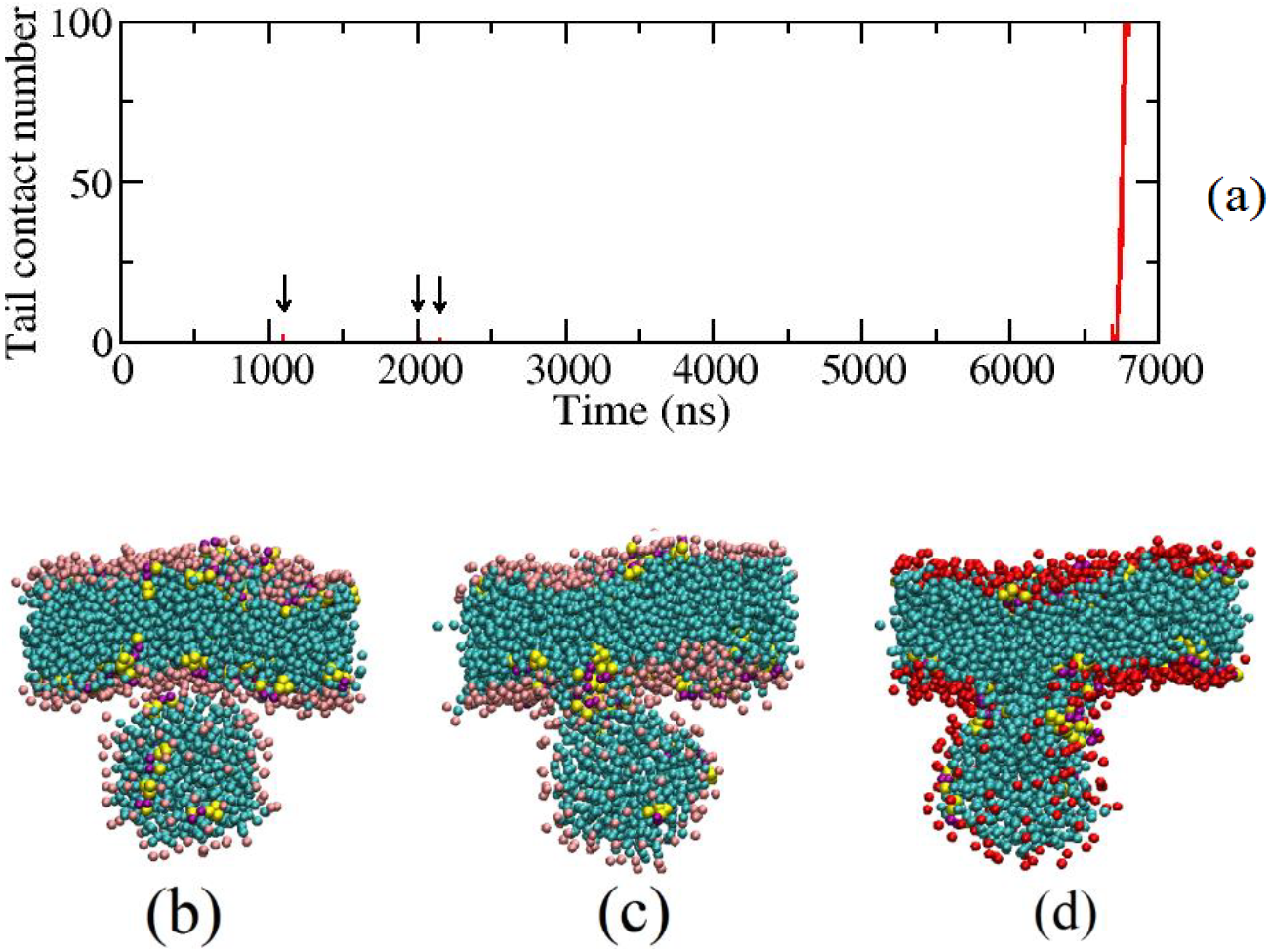
(a) The total contact number, *N*_*t*_, between the hydrophobic tails of the OA molecules from the micelle and the membrane, as a function of time. The three instances of brief contacts between tails before the formation of a fusion stalk are marked with arrows. (b) Initial adsorption of the micelle to the membrane. (c) A snapshot of the micelle-membrane complex at the beginning of fusion that has a cluster of Phe-Leu dipeptides present in the contact region. (d) A snapshot of a stalk structure following the initial fusion events. Stalk formation is the irreversible step in fusion. Red/pink: head groups of OA; cyan: hydrophobic tails of OA; yellow: dipeptides. Water and ions are not shown for clarity.

The stalk structure identified in the simulations has long been postulated as the intermediate state during membrane-membrane fusion (38–40). In particular, a theoretical model based on elastic theory proposed for fusion involves this structure as a crucial intermediate (67, 68). Although direct experimental confirmation of this structure is difficult due to its transient nature, it has been observed between two membranes under specific dehydration conditions in X-ray diffraction experiments (69). Our MD simulation studies provide a direct computational validation of this crucial state in fusion process.

Although mixing of OA molecules from the micelle and the membrane continues rapidly after stalk formation, eventually leading to the complete integration of the micelle into the vesicle, we have not investigated this process beyond the stalk formation step. Since all OA molecules in a micelle are deprotonated, half of them have to become protonated once incorporated to the membrane in order to restore the equilibrium protonation state of a vesicle at near-neutral pH. This can be captured in constant pH MD (CpHMD) simulations, but such simulations in combination with MARTINI force field do not appear to be very successful. CpHMD simulations of small aggregations of OA molecules (70) and OA membrane (71) in which the shifted cutoff method for electrostatic interactions was used were found to be unsuitable for systems involving high content of ions and protonable molecules. Specifically, the area per OA head-group was calculated to be equal to 26 Å^2^ (71), which was significantly lower than the measured value of 33 Å^2^ (52). The implementation of PME in CpHMD is also problematic, as PME implicitly enforces a uniformly charged background, which has been found to introduce artifacts for systems similar to those of interest here (72).

Although full fusion is completed only when the micelle becomes totally integrated with the membrane, terminating simulations shortly after stalk formation is less restrictive than it might initially appear. This is because stalk formation is anticipated to be the only rate limiting step during fusion between micelles and vesicles, and once it is formed fusion usually proceeds to completion. In contrast, fusion between vesicles involves additional, subsequent barrier associated with the formation of a pore to exchange intra-vesicle contents (48, 73, 74).

### Pathway for stalk formation: hydrophobic patch induced by dipeptides in the contact area

In all simulations, stalk formation was found to correlate with the presence of dipeptides in the contact region (see Fig. 1c), suggesting their direct role in mediating the process. Before fusion, a layer of polar or charged species, such as OA head groups, water molecules and ions, separates hydrophobic tails of lipids from the micelle and the membrane. At the same time, the membrane-embedded hydrophobic peptides diffuse in and out of the contact region and occasionally several of them cluster. For quantitative analysis of aggregation, a dipeptide is considered to be in the vesicle-micelle contact region if a bead from its hydrophobic side chain is within the VDW distance from at least one tail bead of lipids from both the micelle and the membrane. A cluster is defined as a group of dipeptides in which each member has at least one bead (group) within the VDW distance from at least one bead of another member. In Fig. S4, the number of Phe-Leu dipeptides in the largest cluster fully located in the contact region is plotted as a function of time. Single Phe-Leu dipeptides or transient clusters consisting of up to 6 molecules are present in the contact region during 10% of the simulation trajectory. A dipeptide can join or leave a cluster on a very fast time scale of a few to tens of *ns*. In the case discussed above, the jump in the cluster size at t = 6.7 *μs* correlates with the stalk formation. The appearance of a larger than average clusters at *t* =1.2, 2, and 2.2 *μs* coincides with a non-zero value of the lipid tail contact parameter, *N*_*t*_, as shown in Fig. 1a, further demonstrating the key role of dipeptides in the stalk formation.

Since the side chains of both amino acids in all dipeptides studied here are hydrophobic, their clustering creates a hydrophobic patch that excludes charged or polar molecules from the contact region. This, in turn, reduces the electrostatic energy barrier to mixing of hydrophobic tails of OA molecules from the membrane and the micelle. A series of snapshots along fusion pathway is shown in Fig. 2. Initially, the interfacial layer is largely hydrophilic (Fig. 2a). Once a cluster of Phe-Leu dipeptide forms in the contact area (Fig. 2b), a sizable hydrophobic patch develops in the same region (Fig. 2c) making lipid tails from both the micelle and the membrane more likely to interact and form a stalk (Fig. 2d). The significance of hydrophobic contacts was recognized earlier in a study in which a reaction coordinate based on this quantity was used (37).

**Figure 2.**
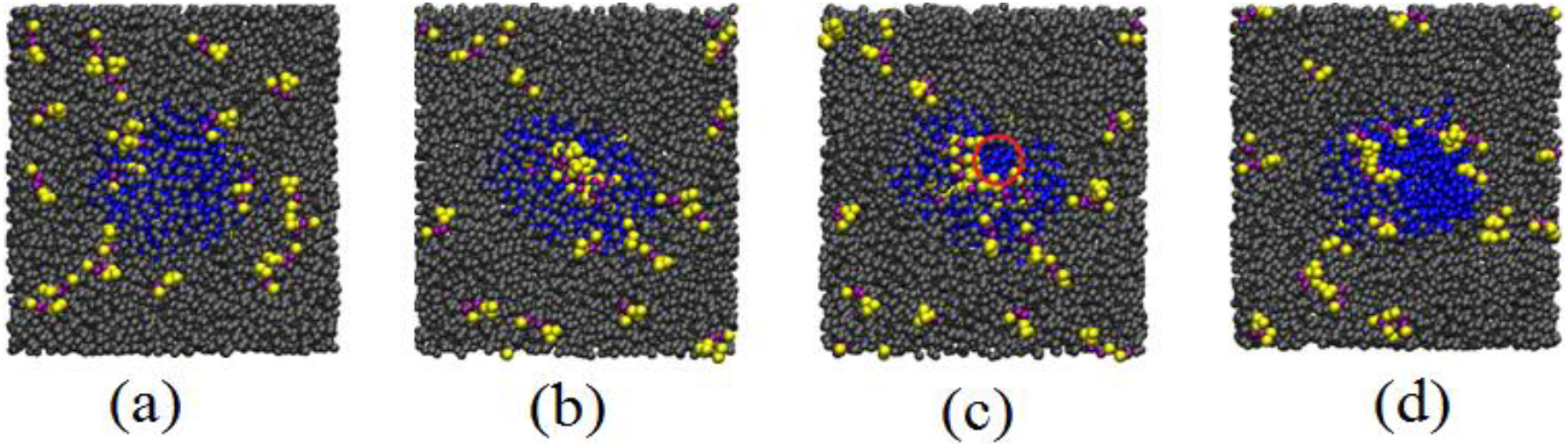
Snapshots (top view along the z-coordinate) along a fusion event in the Phe-Leu system. (a) A layer of charged and polar species between the micelle and the membrane. (b) A cluster of Phe-Leu dipeptides in the contact region. (c) Subsequently formed hydrophobic patch (red circle). (d) The formation of a stalk structure. Black: all charged or hydrophilic species or groups, i.e. water, ions, and head groups of OA molecules; yellow and maroon: side chains and the backbone of Phe-Leu dipeptides, respectively; blue: hydrophobic tails of OA molecules in the micelle. Note the absence of black beads in the hydrophobic patch.

In quantitative analysis, the surface area, *σ*, of the hydrophobic patch at the membrane surface was tracked during the simulation. To calculate *σ*, the contact region at the membrane surface plane was discretized on a small grid (0.1 Å × 0.1 Å). A grid point was considered hydrophilic if it was occupied by a surface hydrophilic or polar molecule or bead. Otherwise it was considered hydrophobic. A hydrophobic patch was identified by way of joining neighboring hydrophobic grid points, and its size was determined by the number of grid points forming the patch. The area of the largest patch defined *σ*, whereas other small, isolated patches were neglected. Plotted in Fig. 3a is the calculated *σ* during 60 ns, directly preceding the fusion. As can be seen, a hydrophobic patch of size *σ* ~ 100 Å^2^ formed around *t* ~ 6690 ns. This patch was sufficient to accommodate four hydrophobic beads with a radius of 2.63 Å. This correlates closely with the formation of contacts between the hydrophobic tails from the micelle and the membrane, as shown in Fig. 3b. However, the hydrophobic patch and the contact of lipid tails were both transient. They lasted for only a few ns, but reappeared later (*t* = 6716 *ns*), this time leading to a successful fusion event, as illustrated by the abrupt, significant increase in both the area of the patch and the contact number. Examination of configurations along the MD trajectory confirms that a stalk is formed at this stage.

**Figure 3.**
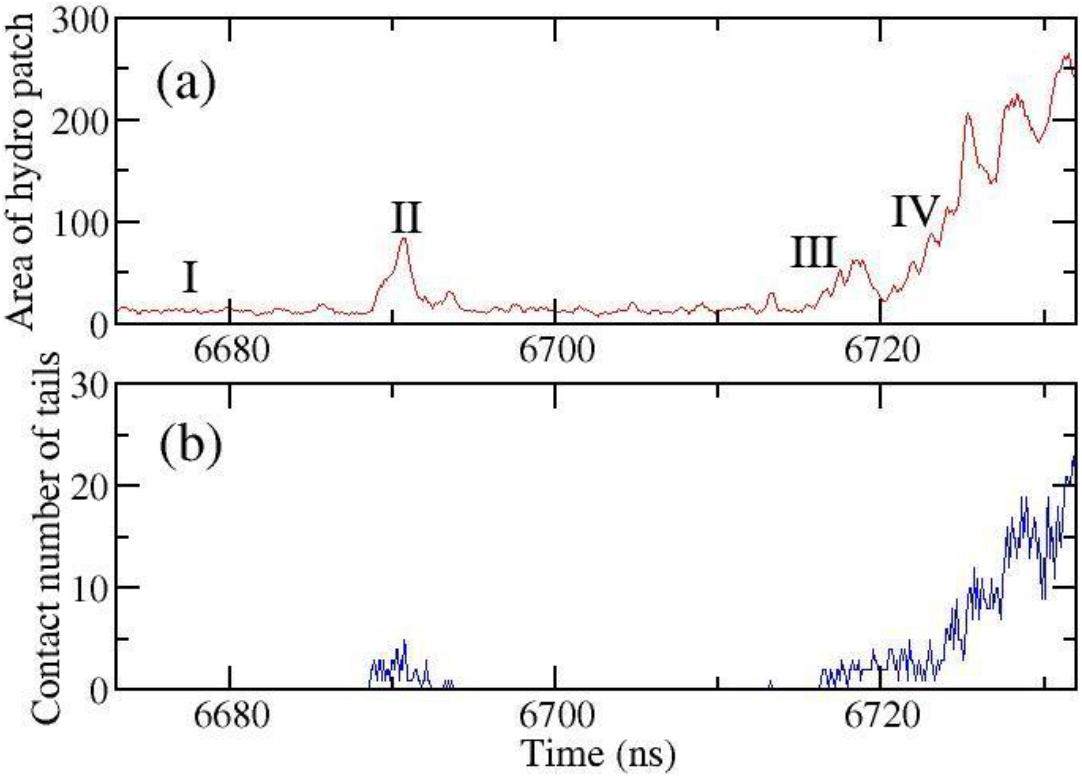
(a) Surface area of the largest hydrophobic patch in the contact region between the micelle and the OA membrane (unit: Å^2^), as a function time. (b) Contact number between hydrophobic tails as a function of time. The data was obtained from one of the Phe-Leu systems. Only a period of time close to the fusion is shown.

Similar involvement of dipeptides in the formation of a fusion stalk was found in simulations of dipeptide-mediated fusion for all dipeptides studied here. The time, *τ*_*s*_, to the formation of a stalk, identified by an abrupt, large jump in the number of contacts between lipid tails from the micelle and the membrane, similar to that shown in Fig. 1a, is listed in Table 1 for all four dipeptides and all simulation sets. The average *τ*_*s*_ is equal to 2.8, 3.6, 7.4, and 6.1 *μs* for the Phe-Phe, Phe-Leu, Val-Val, and Leu-Leu system, respectively. For the first three systems, *τ*_*s*_ obtained from 8 independent simulations was fitted to the cumulative distribution function of finding fusion event at *τ* ≤ *τ*_*s*_ equaling to 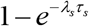, where *τ*_*s*_ is the rate, to ascertain whether the distribution of *τ*_*s*_ follows Poisson statistics. A similar procedure was previously used in analyses of channel-mediated ion crossing events through membranes (72). As can be seen in Fig. S5, the observed distribution of *τ*_*s*_ and the predictions based on the Poisson distribution agree. Even though the number of fusion events is not sufficient to draw firm conclusions, there is no statistical basis to reject the hypothesis that fusion is a Poisson process.

**Table 1.** The time to the fusion stalk formation for all simulation sets for each dipeptide.

| Dipeptides | Number of simulation sets | Time to the stalk formation from each individual simulation ($\mu s$) |
| --- | --- | --- |
| Phe-Phe | 8 | 2.6; 2.1; 5.3; 1.7; 2.4; 0.4; 7.9 |
| Phe-Leu | 8 | 6.7; 3.2; 1.8; 1.2; 1.3; 1.2; 12.2; 1.8 |
| Val-Val | 8 | 19.4; 5.0; 4.4; 10.2; 2.3; 7.1; 10.4; 0.5 |
| Leu-Leu | 4 | 2.3; 12.3; 4.2; 3.0 |

The rate of stalk formation depends on the hydrophobicity of a dipeptide. It is the slowest for Val-Val, which has the smallest hydrophobic side chains among the four dipeptides considered here, whereas the fastest rate is found for Phe-Phe, which has the largest hydrophobic side chain. A natural explanation is that bulkier side chains form larger hydrophobic patches. As in the case of Phe-Leu, the surface area, *σ*, plotted in Fig. S6a as a function of time for representative simulations of each system, always exhibits a sharp increase just before stalk formation, but fluctuations in this parameter persist even before successful fusion. To illuminate further the connection between the formation of large hydrophobic patches and the hydrophobicity of the dipeptides, we plot in Fig. S6b the histograms of *σ* in the range > 20 Å^2^, normalized to the peak distribution at *σ* = 3.5 Å^2^. To obtain the histogram for each dipeptide, data from all simulation sets have been combined. The largest values of *σ* are found for Phe-Phe and the smallest for Val-Val, consistent with the dependence of the fusion rate on peptide hydrophobicity. For comparison, the same histogram has been also calculated for the pure OA membrane-micelle system. In this system, there are practically no hydrophobic patches beyond 25 Å^2^, even though MD simulations extended for 90 *μs*, which is 2-4 times longer than the combined simulations in the presence of a dipeptide. This further illustrates the role of hydrophobic dipeptides in facilitating micelle-vesicle fusion.

It has been previously suggested that the initial contact between hydrophobic tails is the key pre-stalk transition state structure for membrane-membrane or vesicle-vesicle fusion, although observing this structure in computer simulations required either pre-induced splayed tails (38, 39), inducing a stalk by applying an external field (76), or cross-linkage between opposed membranes (40). Hydrophobic peptides facilitate the formation of the fusion stalk to the extent that it can be directly observed in MD simulations.

### Transition state for stalk formation and committor probability calculations

On the basis of the results discussed in the previous section, it is reasonable to propose that *σ* is a good reaction coordinate to describe vesicle/micelle fusion in the presence of dipeptides independently of their specific sequence. Here we demonstrate that this is indeed the case. Further, we identify the transition state for stalk formation along *σ* and estimate the free energy barrier associated with this process.

It is often considered that the ideal reaction coordinate is committor probability. For a given configuration, it is defined as the fraction of trajectories initiated from this configuration that reaches the product state rather than the reactant state first (77). In the transition state, the committor probability is equal to 0.5 (78, 79). Although the committor probability is the best formal choice for the reaction coordinate, determining and interpreting complex surfaces of a constant committor probability might be quite difficult. For this reason, it is often more practical and informative to use a reaction coordinate that is simpler but remains a good approximation to the committor probability. For such coordinate, the committor probability at a fixed value of the coordinate is no longer constant, but instead forms a narrow distribution and its average value increases monotonically between the reactant and product states. If this is not the case the selected reaction coordinate may not be suitable for the process of interest.

To calculate the committor probability along *σ*, 21 and 22 configurations near the fusion stalk formation were selected from the long MD trajectories for Phe-Leu and Phe-Phe, respectively. For Val-Val and Leu-Leu, six configurations were chosen. As examples, four configurations of the Phe-Leu system labeled as I, II, III, and IV that were selected from the simulation trajectory near the beginning of fusion are shown in Fig. S7. The time points corresponding to these configurations are marked in Fig. 3. They correspond, respectively, to a state that precedes the formation of the first, sizable hydrophobic patch, a failed fusion attempt, the beginning of successful fusion, and initial mixing of hydrophobic tails from the micelle and the vesicle during stalk formation.

To calculate committor probabilities, forty MD trajectories initiated from each of the selected configurations were obtained. For each trajectory, initial velocities of all beads were sampled from the Maxwell-Boltzmann distribution. Each trajectory was terminated when either the stalk was formed, or the hydrophobic patch was reduced to less than 5 Å^2^. These two end states correspond, respectively, to the product formation and the return to the reactant state. The fraction of trajectories that ended forming the stalk provided the estimate of the committor probability for a given configuration. The average committor probabilities for configurations I to IV in Fig. S7 were 0, 0.7, 0.5, and 1, with an estimated statistical error of 0.15, calculated as 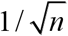 for *n* = 40 independent trajectories. In configuration III, which is very close to the transition state, the hydrophobic tail of one OA molecule from the micelle (red) is in contact with that of an OA molecule from the membrane (blue) through a cluster of dipeptides. As we have already pointed out, this feature is required for stalk formation, but not every configuration exhibiting this feature is on the path to a successful fusion event. In configuration II, which is also close to the transition state, hydrophobic tails from the micelle and the membrane are in contact, but this configuration is not on a successful fusion path in the long MD simulation (see Fig. 3). Instead, it is a part of an earlier, failed fusion attempt. This is a result of an unfavorable distribution of particle velocities, as 70% of velocities sampled from the Maxwell-Boltzmann distribution for this configuration lead to fusion.

The calculated committor probability as a function of *σ* for all systems with the dipeptides is shown in Fig. 4. These two variables are highly correlated with only modest dispersion. For linear fit the correlation coefficient is 0.875. For a slightly nonlinear fit to a function (1 − exp(−*αx*)) / (1+ exp(*α*x)) with the optimized value of the fitted parameter *α* equal to 0.019, the correlation coefficient increases to 0.950. No statistically significant dependence on the type of dipeptide was found, indicating that *σ* is a universally good approximation to the exact reaction coordinate for all systems studied here. The quality of a reaction coordinate can be characterized more accurately through the histogram test (78, 79). On this basis, it is possible to calculate measures that quantify reaction coordinate errors (80). This approach was not pursued here because the number of trajectories was not sufficient for its reliable implementation.

**Figure 4.**
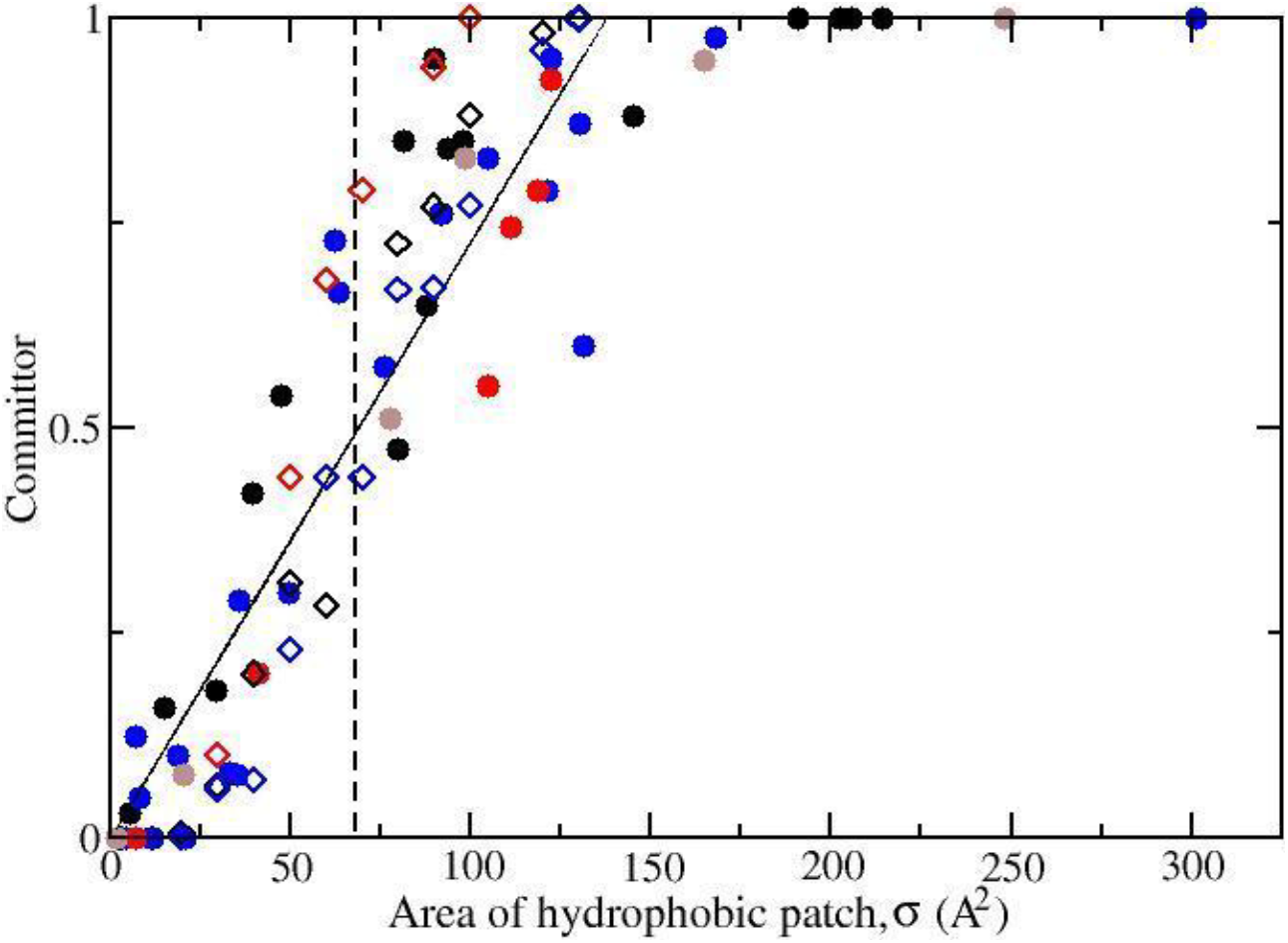
The committor probability for the fusion stalk formation as a function of the area of the hydrophobic patch in the contact region. Solid black, red, blue and brown circle symbol is for Phe-Leu, Val-Val, Phe-Phe, and Leu-Leu system, respectively (from MD simulations). Hollow diamond symbol is for committor probabilities obtained from 1D diffusion trajectories generated from Monte Carlo simulations on the free energy profile for fusion (see next section for free energy calculation), with black, red, and blue for Phe-Leu, Val-Val, and Phe-Phe. The transition state (committor probability = 0.5) is located at *σ* approximately equal to 68 Å^2^, which is marked by the dashed line.

### Free energy barrier to fusion

Once a good reaction coordinate has been identified, the free energy along this coordinate can be determined. A host of methods based on equilibrium statistical mechanics are available for this purpose (81). Unfortunately, these methods cannot be readily applied to this problem. Although it is possible to calculate the value of *σ* for every configuration, no technique for statistical sampling of configurations at a fixed value or a predefined range of *σ* is available.

Alternatively, the free energy as a function of *σ*, *A(σ)*, can be extracted from non-equilibrium simulations assuming the validity of the diffusion model with steady state condition:

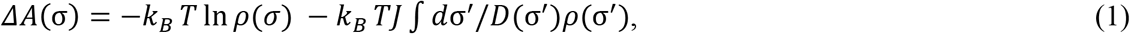

where *k*_*B*_ is the Boltzmann constant, *T* is temperature, *J* is the steady state flux, and *ρ(σ)* and *D(σ)* are position-dependent population density and diffusion coefficient, respectively. This formula also cannot be applied directly because it requires the knowledge of the diffusion coefficient.

Without prior knowledge of *A(σ)*, the method of choice for determinin*D*g *(σ)* is based on the fluctuation-dissipation theorem and involves calculating the force-force correlation function at fixed *σ*(82). However, as mentioned before, it is not clear how to hold *σ* fixed. Another approach is to extract *A(σ)* and *D(σ)* from simulations simultaneously (83). This method has a disadvantage that all inaccuracies of the model and statistical imprecision of the data are captured in *D(σ)*. As a consequence, the diffusion coefficient determined by way of this approach might deviate significantly from the real *D(σ)*.

To circumvent these difficulties, we used a method based on mean first-passage time (MFPT) proposed by Wedekind and Reguera (84). Here, MFPT at *σ* is defined as the time needed for a MD trajectory initiated at 0 to reach*σ*, averaged over all trajectories considered. In this method, *A(σ)*, can be reconstructed from non-equilibrium, steady-state MD simulations without calculating the diffusion coefficient:

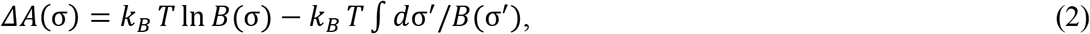

where *B(σ)* is a function that depends on the position-dependent population density*ρ(σ)* and the MFPT, *τ*(*σ)*:

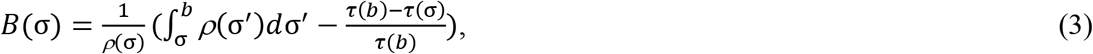

with adsorbing boundary condition applied at *σ* = *b*, which equals to 135 Å^2^, at which the committor probability is equal to 1 for the fusion process (obtained from the linear fitting of committor probability to *σ*, as plotted in Fig. 4).

Before applying to the fusion process, the approach was validated in a benchmark case in which diffusive trajectories initiated in one potential well and absorbed in the other were generated by way of a stochastic Monte Carlo (MC) method. The free energy profile reconstructed by way of Eqs. 2 and 3 was found to agree with the input potential.

A straightforward approach to calculating *τ(σ)* is to generate an ensemble of trajectories that typically are terminated when the system either reaches the product or returns to the reactant state. For each trajectory, the time needed to reach different values of *σ* for the first time is recorded and the average over all trajectories yields *τ(σ)*. To obtain such trajectories, we used 8 long fusion trajectories previously obtained for the Phe-Leu, Phe-Phe, and Val-Val systems. Each of these trajectories was divided to a number of shorter trajectories. Every time the system returned to the reactant state in which the size of the hydrophobic patch decreased below 5 Å^2^, a trajectory was terminated. This was also the beginning of the next trajectory. This new trajectory could be considered as statistically independent of the previous one if the decorrelation time was markedly shorter than the MFPT, an approximation that appeared to hold satisfactorily. A disadvantage of this simple approach is that the number of trajectories decreases markedly with increasing *σ* (see Fig. 5b). As a result, the statistics for large *σ* suffers and, therefore, the reconstructed *A(σ)* is approximate.

**Figure 5.**
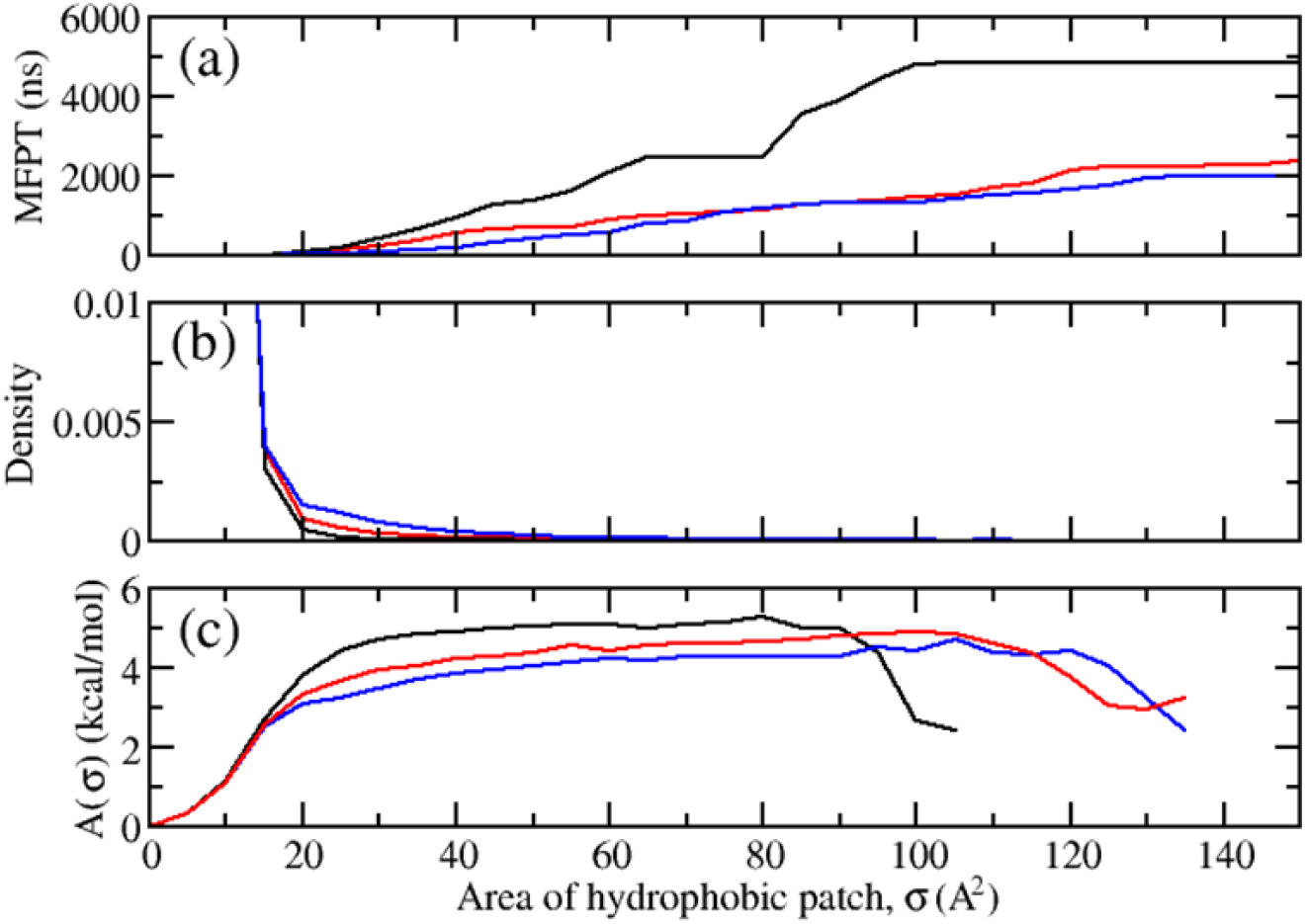
(a) The mean first-passage time (MPFT) as a function of its size, σ, in the contact region. (b) Normalized population density for *σ*. (c) The free energy profiles for the formation of a fusion stalk as a function of σ. Black, red, and blue is for the Val-Val, Phe-Leu, and Phe-Phe system, respectively.

In the presence of Val-Val, which is the least hydrophobic of the three dipeptides considered here, the MFPT is longer and the free energy barrier to fusion of approximately 5 kcal/mol is higher than in the presence of the other two dipeptides by 0.5-1 kcal/mol (see Fig. 5a and 5c, respectively).

If the calculated free energy profiles are used in the diffusion model then, assuming the same diffusion constant for all three systems, the estimated fusion rate in the Val-Val system is 1.6 and 2.5 times slower than the fusion rates for the Phe-Leu and Phe-Phe systems, respectively. This is in good agreement with the corresponding ratios calculated directly from the average lengths of reactive fusion trajectories, which are 2.0 and 2.6, indicating that both methods yield consistent results. This consistency is further confirmed by observing that committor probabilities calculated from MD simulations and from the diffusion model with the reconstructed free energy profiles are very similar (see Fig. 4).

The energy barrier for the formation of the stalk structure has been previously estimated at 90 kcal/mol or more (85, 86) on the basis of membrane elastic theory for membrane-membrane fusion (67, 68). A markedly lower value of 24 kcal/mol was estimated from the measured rate of opening and closing of exocytotic fusion pore (42). A modified stalk structure with tiled lipid tails was later considered to reduce the unrealistically high barrier (87, 88). It is noted that the stalk structure assumed in the elastic theory model may not be accurate and the electrostatics repulsion from the hydrophilic layer at the membrane surface is neglected. Free energy profiles for the membrane-membrane fusion were also calculated in MD simulations (49, 74), in which a guiding force was applied along a pathway in which the stalk was assumed as the intermediate. These calculations yielded a barrier of ~ 24 – 50 kcal/mol for two opposed DOPC/DOPE, DMPC/POPC, and DMPC/DOPE membranes (74, 89). As shown in our study, the presence of hydrophobic dipeptides drastically lowers the energy barrier to only 4 - 5 kcal/mol.

### Accelerated flip-flop of OA in the membrane

Although the flip-flop motion of OA molecules across the OA membrane is not directly related to the formation of the fusion stalk, it plays a key role in the equilibration of OA molecules absorbed from the micelle to the outer leaflet of the membrane bilayer. Flip-flop is the main mechanism to balance the number of OA molecules between the outer and the inner leaflet. Plotted in Fig. S8 is the cumulative number of OA flip-flops in the presence of different dipeptides as a function of time. The flip-flop rate of 2.5 to 3 *μs*^−1^ is approximately 50% faster than the flip-flop rate previously calculated for the pure OA membrane (90). This change can be attributed to the reduced structural order in the tails of OA molecules in the presence of dipeptides, which lowers the free energy barrier to the flip-flop motion. This means that hydrophobic dipeptides not only promote stalk formation, which is the first step in fusion, but also accelerate the subsequent step.

## Conclusion

It was previously determined experimentally that the presence of hydrophobic dipeptides promotes fusion between vesicles and micelles (24). In this study, the underlying mechanism of this process is examined. Analogously to vesicle-vesicle fusion, micelles fuse with vesicles through the formation of a stalk joining these two structures, whereby hydrophobic groups from the vesicle and the micelle make direct contact. This initiates irreversible mixing that results in vesicle growth. In the absence of dipeptides, fusion is slow, as hydrophobic groups in vesicles and micelles are separated from each other by hydrophilic headgroups, water molecules and ions.

If hydrophobic dipeptides, such as Phe-Phe, Leu-Leu, Val-Val, and Phe-Leu, are present in vesicles they tend to accumulate at the vesicle surfaces forming intermittent, dynamic patches. The clustering of the hydrophobic side chains provides a hydrophobic core region in the contact region between vesicles and micelles. Hydrophilic groups, water and ions are excluded from this region. This, in turn, facilitates direct interactions between hydrophobic tails from vesicles and micelles and lowers the free energy barrier to stalk formation that further leads to fusion. For the dipeptides studied here, this free energy barrier has been estimated at 4-5 kcal/mol and decreases with increased hydrophobicity of the dipeptide side chains. It appears that the size of a hydrophobic patch in the contact area is a good reaction coordinate since it correlates nearly linearly with the probability of stalk formation. Accelerated flip-flop of OA molecules from micelles incorporated into vesicles induced by the embedded dipeptide additionally facilitates fusion.

Since the mechanism of vesicle growth described here is general, other small peptides and hydrophobic molecules that tend to accumulate at the water-membrane interface (91) might mediate the same process. The same mechanism might also be at play in vesicle-vesicle fusion. In the context of the origin of life, growth of vesicles capable of synthesizing interfacially active peptides and metabolites in their interior was more facile than growth of vesicles that did not have these capabilities, thus providing a universal, evolutionary mechanism of coupling metabolism and compartmentalization.

## Author Contributions

C.W. and A.P. planned the research. C.W. conducted the MD simulations. Both authors analyzed the data, performed the research, and wrote the article.

## Acknowledgments

Support for this research was provided by NASA’s Planetary Science Division Research Program. All simulations were performed at the NASA Advanced Supercomputing (NAS) Division.

